# Phase separation-aided compartmentalization of protein-protein interactions in cells

**DOI:** 10.1101/860817

**Authors:** Hongrui Liu, Jing Wang, Pilong Li

## Abstract

Protein-protein interactions (PPIs) are of great significance in many biological activities and diseases. It is essential to distinguish between direct, indirect and false PPIs in order to accurately understand their biological functions. Here we report a method based on <u>p</u>hase separation-<u>a</u>ided <u>c</u>ompartmentalization of <u>p</u>rotein-protein interactions in <u>c</u>ells (PACPIC), which is tailored for identification of direct and indirect PPIs in cells, using recruitment as the readout.

## Introduction

PPIs are vital to all cellular activities including gene expression, signal transduction, proliferation and apoptosis. Aberrant PPIs are responsible for many diseases such as neurodegenerative diseases and cancers. By studying PPIs, we can understand the functions of specific proteins within biological networks and screen for potential drugs to treat diseases. Numerous approaches, such as co-immunoprecipitation, yeast two-hybrid assays and phage display, by means of biophysics, biochemistry, or genetics, have been applied to investigate PPIs in cells (Zhou et al., 2016). However, it is still challenging to identify direct, indirect, or false PPIs.

Cells contain numerous membraneless organelles that compartmentalize biochemical reactions. Emerging evidence indicates that these compartments are assembled via liquid-liquid phase separation (LLPS) driven by multivalent interactions of scaffolding constituents (Banani, Salman F. et al., 2017). Other constituents or clients partition into these organelles via specific interactions (Banani, Salman F. et al., 2016). Inspired by the architecture of cellular membraneless organelles (Watanabe, Taku et al., 2017; Nakamura, Hideki et al., 2018; Zhang, Qiang et al., 2018; Nakamura et al., 2019), we established a method based on phase separation—PACPIC—for reliably investigating direct and indirect interactions between proteins that are known and/or suspected to form a complex in cells, with recruitment as the readout. The design principle of PACPIC is as follows: 1) a system capable of robustly undergoing phase separation (scaffold) is used to form membraneless compartments in cells; 2) one component of the PPI of interest is fused with the system and hence is enriched in the compartments; 3) the recruitment of the other component(s) (client) into the compartments serves as the readout for the interaction (Fig. 1a,b).

**Fig. 1.**
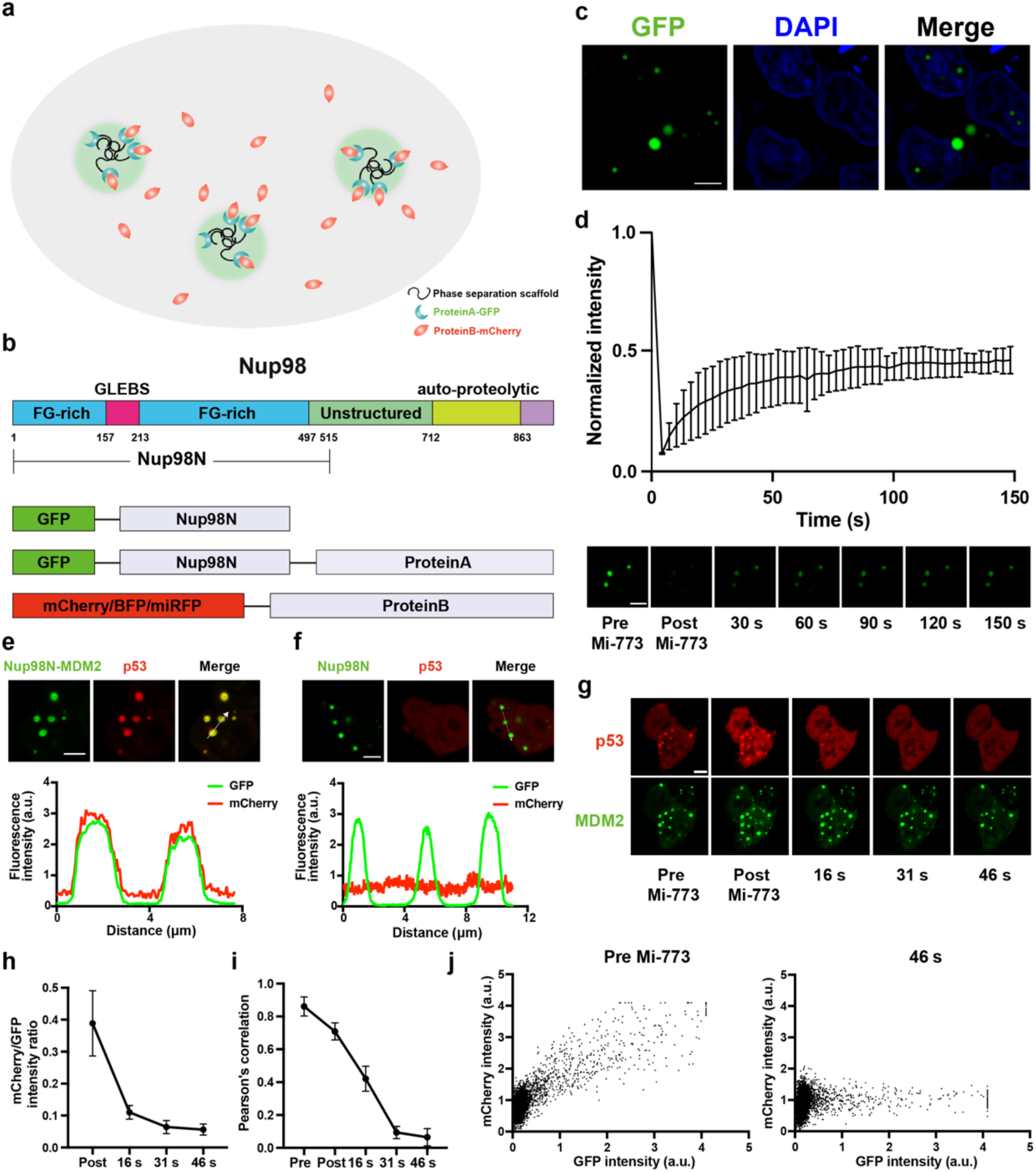
Establishment of a robust phase separation system in cells. **a**, Schematic diagram showing a co-transfection experiment with two plasmids. One plasmid encodes protein A fused to a GFP-tagged phase separation-prone scaffold protein and the other encodes protein B fused to mCherry. In cells co-transfected with both plasmids, protein A is enriched within green fluorescent puncta, which are phase-separated compartments formed by the phase separation-prone scaffold protein. If protein A and protein B directly interact, protein B will be recruited into the phase-separated compartments by binding to protein A. Hence, the red fluorescence signal is also enriched in the GFP-labeled compartments, which serves as the readout for the interaction. **b**, The domain structure of human Nup98. The FG-rich region at the N terminus of Nup98, also known as the FG-domain, is prone to phase separation. The structures of GFP-Nup98N, GFP-Nup98N-protein A and mCherry/BFP/miRFP-protein B are shown below. **c**, Confocal images of HEK 293 cells overexpressing GFP-Nup98N, with the nucleus stained by DAPI. GFP-Nup98N forms green spherical droplets in both the nucleus and cytoplasm of the cells. **d**, Plot of normalized GFP fluorescence intensity of GFP-Nup98N puncta versus time after total photobleaching. Confocal images of GFP-Nup98N at different time points are shown below. The green fluorescence signal started to recover immediately after bleaching by a 488-nm wavelength laser, and up to 50% of the original fluorescence intensity was recovered within 150 seconds. **e**, Confocal images of co-transfection of GFP-Nup98N-MDM2 and mCherry-p53 in cells, with line analysis of fluorescence intensity below. The p53 fusion protein, as indicated by the mCherry signal, was recruited to the green droplets formed by GFP-Nup98N-MDM2. The result was confirmed by plotting the fluorescence intensities of GFP and mCherry along the line indicated by the white arrow in the merged confocal. **f**, Confocal images of co-transfection of GFP-Nup98N and mCherry-p53 in cells, with line analysis of fluorescence intensity below. The mCherry signal remained evenly distributed when co-transfected with GFP-Nup98N. The result was confirmed by plotting the fluorescence intensities along the line indicated by the white arrow in the merged confocal image. **g-j**, Treatment of cells expressing GFP-Nup98N-MDM2 and mCherry-p53 with Mi-773, an inhibitor of the MDM2/p53 interaction. **g**, Confocal images of the time-course of inhibitor treatment. The mCherry signal within the phase-separated puncta started to weaken immediately after the addition of 5 μM Mi-773, and very little mCherry was enriched within the puncta at 46 seconds after treatment. **h**, Plot of the ratio of mCherry/GFP intensity within puncta versus time. The mCherry intensity within puncta decreased dramatically relative to GFP after treatment with Mi-773. **i**, Plot of Pearson’s correlation coefficient for the intensity of mCherry and GFP versus time. The correlation between the two intensities decreased significantly with time and was close to 0.1 at 46 seconds after treatment. **j**, Scatter diagrams of the intensity of mCherry versus GFP before inhibitor treatment and at 46 seconds after treatment. The mCherry intensity correlated significantly with the GFP intensity before treatment, while there was barely any correlation between the two at 46 seconds after treatment. Nup98N, N-terminus of Nup98. n = 3. Values are means ± standard deviation. All scale bars, 5 μm.

## Results

### Establishment of a robust protein scaffold for phase separation in cells

Specifically, we fused the N-terminal half of Nup98 (Nup98N, a.a. 1-515) with green fluorescent protein (GFP) to generate GFP-Nup98N (Fig. 1b). Nup98N readily undergoes phase separation *in vitro* and in cells (Banani, Salman F. et al., 2017; Frey, Steffen, and Dirk Görlich, 2007; Schmidt, Hermann Broder, and Dirk Görlich, 2015). GFP-Nup98N formed green micron-scale membraneless compartments upon overexpression in HEK 293 cells (Fig. 1c). Fluorescence recovery after photobleaching experiments showed that the GFP-Nup98N molecules were dynamic as about half of the bleached GFP signal recovered within 150 seconds after bleaching (Fig. 1d). Based on these results, we concluded that GFP-Nup98N is a robust scaffold for driving compartmentalization via phase separation.

### Nup98N-mediated compartments support client PPI-mediated recruitment

To test whether Nup98N-mediated compartments support client PPI-mediated recruitment, we fused the p53 transactivation helix (abbreviated to p53 hereafter) to the C-terminus of GFP-Nup98N, which yielded GFP-Nup98N-p53. When GFP-Nup98N-p53 was co-expressed with mCherry-labeled MDM2 (mCherry-MDM2) in cells, the mCherry fluorescence signal was enriched within the green compartments (Fig. 1e), presumably due to the well-known MDM2/p53 interaction (Momand, Jamil et al., 1992). The enrichment is p53-dependent as mCherry-MDM2 was not recruited into GFP-Nup98N-mediated compartments (Fig. 1f). To further confirm that the enrichment of red fluorescence signal was due to the interaction between MDM2 and p53, we treated the cells expressing GFP-Nup98N-p53 and mCherry-MDM2 with Mi-773, a potent inhibitor of the MDM2/p53 interaction. Immediately after adding Mi-773 into the medium, the mCherry signal within the puncta started to decrease and little extra mCherry remained within puncta 46 seconds after treatment as judged by imagines (Fig. 1g) and the ratios between the intensity of mCherry over that of GFP (Fig. 1h). There was also a clear correlation recession between the overall intensity of mCherry and GFP within the cells, as is shown by the Pearson’s correlation coefficient (Fig. 1i) and the scatter plots of the intensity of mCherry versus GFP before and 46 seconds after treatment (Fig. 1j). These data together indicate that Nup98N can yield robust compartments for recruitment-based PPI detection in cells, thus laying a firm foundation for PACPIC.

### Recruitment of proteins of PRC2 complex into phase-separated compartments via direct interactions

One of the challenges of characterizing PPIs in cells is to discern the specific interactions (direct and indirect) from nonspecific ones. It is even more challenging to distinguish between direct and indirect PPIs. Since PACPIC detects the enrichment of client proteins in membraneless compartments versus a surrounding environment, which is against the concentration gradient and thus has to go over the energy barrier, authentic interactions—direct or indirect—are required for substantial partitioning into compartments. As such, detection of nonspecific PPIs can be minimized. We used the Polycomb repressive complex 2 (PRC2) as an example to test whether PACPIC can distinguish direct PPIs from indirect ones. PRC2 is comprised of four core subunits: Enhancer of zeste 1 or 2 (EZH1/2), Embryonic ectoderm development (EED), Supressor of zeste 12 (SUZ12), and Retinoblastoma-binding protein 4 or 7 (RBBP4/7) (Margueron, Raphaël, and Danny Reinberg, 2011). It is known that RBBP4 directly contacts SUZ12, but not EZH2 or EED (Ciferri, Claudio et al., 2012; Kasinath, Vignesh et al., 2018). We generated three expression vectors encoding SUZ12, EZH2, and EED fused to the C-terminus of GFP-Nup98N. Numerous green puncta formed in cells expressing these vectors (Fig. 2a). In cells co-transfected with mCherry-RBBP4 and GFP-Nup98N-SUZ12, mCherry signal was clearly enriched in the compartments (Fig. 2b), consistent with the direct interaction between SUZ12 and RBBP4. In cells co-transfected with mCherry-RBBP4 and GFP-Nup98N-EZH2 or GFP-Nup98N-EED, the mCherry signals in the compartments were of similar intensity to those in the ambient environment (Fig. 2c,d). This is consistent with the lack of direct interaction between RBBP4 and EZH2 or EED.

**Fig. 2.**
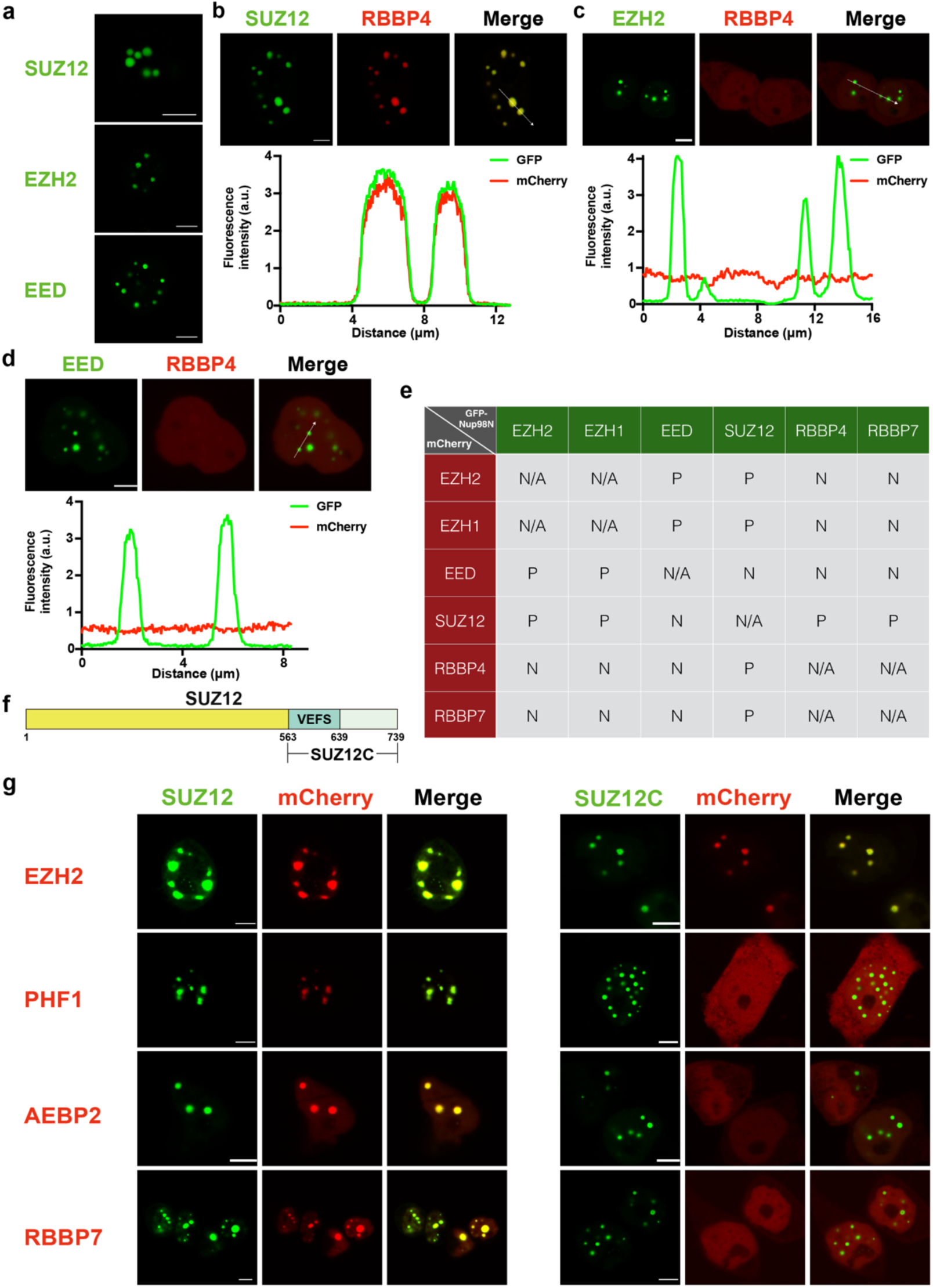
Compartmentalization of direct protein-protein interactions in cells. **a**, Confocal images of HEK 293 cells overexpressing GFP-Nup98N-SUZ12, GFP-Nup98N-EZH2, or GFP-Nup98N-EED. All three images show multiple green spherical puncta within cells. **b**, Confocal images of HEK 293 cells overexpressing GFP-Nup98N-SUZ12 and mCherry-RBBP4, with line analysis of fluorescence intensity below. GFP-Nup98N-SUZ12 formed green spherical puncta within cells and the mCherry signal of the RBBP4 fusion protein was significantly enriched in GFP puncta. The result was confirmed by plotting the fluorescence intensities along the line indicated by the white arrow in the merged confocal image. **c**, Confocal images of HEK 293 cells overexpressing GFP-Nup98N-EZH2 and mCherry-RBBP4, with line analysis of fluorescence intensity below. GFP-Nup98N-EZH2 formed green spherical puncta within cells, while the mCherry signal of RBBP4 remained evenly distributed in cells and failed to be enriched in the GFP puncta. The result was confirmed by plotting the intensity of fluorescence along the line indicated by the white arrow in the merged confocal image. **d**, Confocal images of HEK 293 cells overexpressing GFP-Nup98N-EED and mCherry-RBBP4, with line analysis of fluorescence intensity below. GFP-Nup98N-EED formed green spherical puncta within cells, while the mCherry signal of RBBP4 remained evenly distributed in cells and failed to be enriched in the GFP puncta. The result was confirmed by plotting the fluorescence intensities along the line indicated by the white arrow in the merged confocal image. **e**, Summary table of all pair-wise interactions of the four PRC2 core subunits, EZH1/2, SUZ12, EED, RBBP4/7. The six cDNAs encoding EZH1, EZH2, SUZ12, EED, RBBP4, and RBBP7 were fused to both mCherry and GFP-Nup98N, resulting in six scaffold vectors and six client vectors. P stands for positive, i.e. red fluorescence was enriched within GFP puncta. N stands for negative, i.e. red fluorescence was not enriched within GFP puncta. **f**, The domain structure of human full-length SUZ12 and SUZ12C (C-terminal region of SUZ12). EZH2 binds to SUZ12C, while RBBP7, PHF1 and AEBP2 do not. **g**, Confocal images of pair-wise interactions between different mCherry-fused subunits (EZH2, RBBP7, PHF1 and AEBP2) and GFP-fused SUZ12 (full-length SUZ12) or SUZ12C (C-terminal region of SUZ12). All the mCherry-fused subunits were enriched in compartments containing GFP-SUZ12. Only mCherry-EZH2 was enriched in compartments containing GFP-Nup98N-SUZ12C. The other subunits, which bind to the N-terminal part of SUZ12, were not enriched. All scale bars, 5 μm.

We have now characterized two extreme examples of PACPIC, i.e. direct interaction and indirect interaction. Next, we examined all pair-wise interactions of the four core subunits, EZH1/2, SUZ12, EED, and RBBP4/7. We fused all six cDNAs encoding human EZH1, EZH2, SUZ12, EED, RBBP4, and RBBP7 to GFP-Nup98N and mCherry, resulting in six scaffold vectors and six client vectors. Overall, the pairwise co-expression results supported each other and were consistent with the reported direct interactions and indirect interactions, except for SUZ12 and EED (Fig. 2e). The latter two have direct contact within a recently solved electron microscopy structure of the PRC2 complex (Kasinath, Vignesh et al., 2018). However, we failed to detect the preferential partitioning of one into compartments enriched with the other (Fig. 2e). We speculate that the interaction between SUZ12 and EED might require the presence of EZH2 to maintain the right conformation.

To further demonstrate the robustness and effectiveness of PACPIC in detecting direct PPIs, we made a truncation to remove the region of SUZ12 that binds to some accessory proteins (Fig. 2f). SUZ12 adopts an extended conformation in the PRC2 complex (Ciferri, Claudio et al., 2012; Chen et al., 2018; Kasinath, Vignesh et al., 2018). Multiple PRC2 core subunits and accessory proteins bind SUZ12 via different surfaces. EZH2 binds to the C-terminus of SUZ12, while RBBP7, PHF1, and AEBP2 bind to its N-terminus (Chen et al., 2018; Kasinath, Vignesh et al., 2018; Youmans, Daniel T., Jens C. Schmidt, and Thomas R. Cech., 2018). We made a scaffold construct with the N-terminus of SUZ12 removed, GFP-Nup98N-SUZ12C. In binary co-transfection experiments with full-length SUZ12, there was marked enrichment of EZH2, RBBP7, PHF1, or AEBP2 within GFP-Nup98N-SUZ12 compartments (Fig. 2g, left panels). However, among the above four subunits, EZH2, but not RBBP7, PHF1, or AEBP2, was enriched in GFP-Nup98N-SUZ12C compartments (Fig. 2g, right panels). These results demonstrated the robustness and accuracy of PACPIC.

### Recruitment of proteins of PRC2 complex into phase-separated compartments via indirect interactions

For indirect clients to be enriched in the compartments, bridging proteins are required. As PACPIC relies on overexpressed scaffolds and clients, endogenous bridging proteins, which are often of low abundance, are likely insufficient to recruit substantial amounts of indirect clients to the compartments. If this reasoning is valid, over-expression of bridging proteins should enable enrichment of indirect clients in compartments. To test this hypothesis, we selected two pairs of indirect interaction partners within the PRC2 complex, EZH2/RBBP4 and SUZ12/EED (Fig. 2e). Binary co-transfection experiment using GFP-Nup98N-EZH2 and miRFP-RBBP7 confirmed that the miRFP signal was not enriched in the green EZH2-containing compartments (Fig. 3a). In contrast, a ternary co-transfection experiment using GFP-Nup98N-EZH2, miRFP-RBBP7, and mCherry-SUZ12 resulted in compartments enriched with both miRFP and mCherry signals, consistent with the notion that SUZ12 acts as a bridge to recruit RBBP7 into EZH2-containing compartments (Fig. 3b). Similarly, a binary co-transfection experiment using GFP-Nup98N-SUZ12 and BFP-EED confirmed that the BFP signal was not enriched in green compartments (Fig. 3c). In contrast, a ternary co-transfection experiment using GFP-Nup98N-SUZ12, BFP-EED, and mCherry-EZH2 resulted in compartments enriched with both mCherry and BFP signals, consistent with the notion that EZH2 bridges EED into SUZ12-containing compartments (Fig. 3d). Taken together, these data indicate that indirect interactions can be unambiguously demonstrated by combining binary and ternary co-transfection experiments in the PACPIC system.

**Fig. 3.**
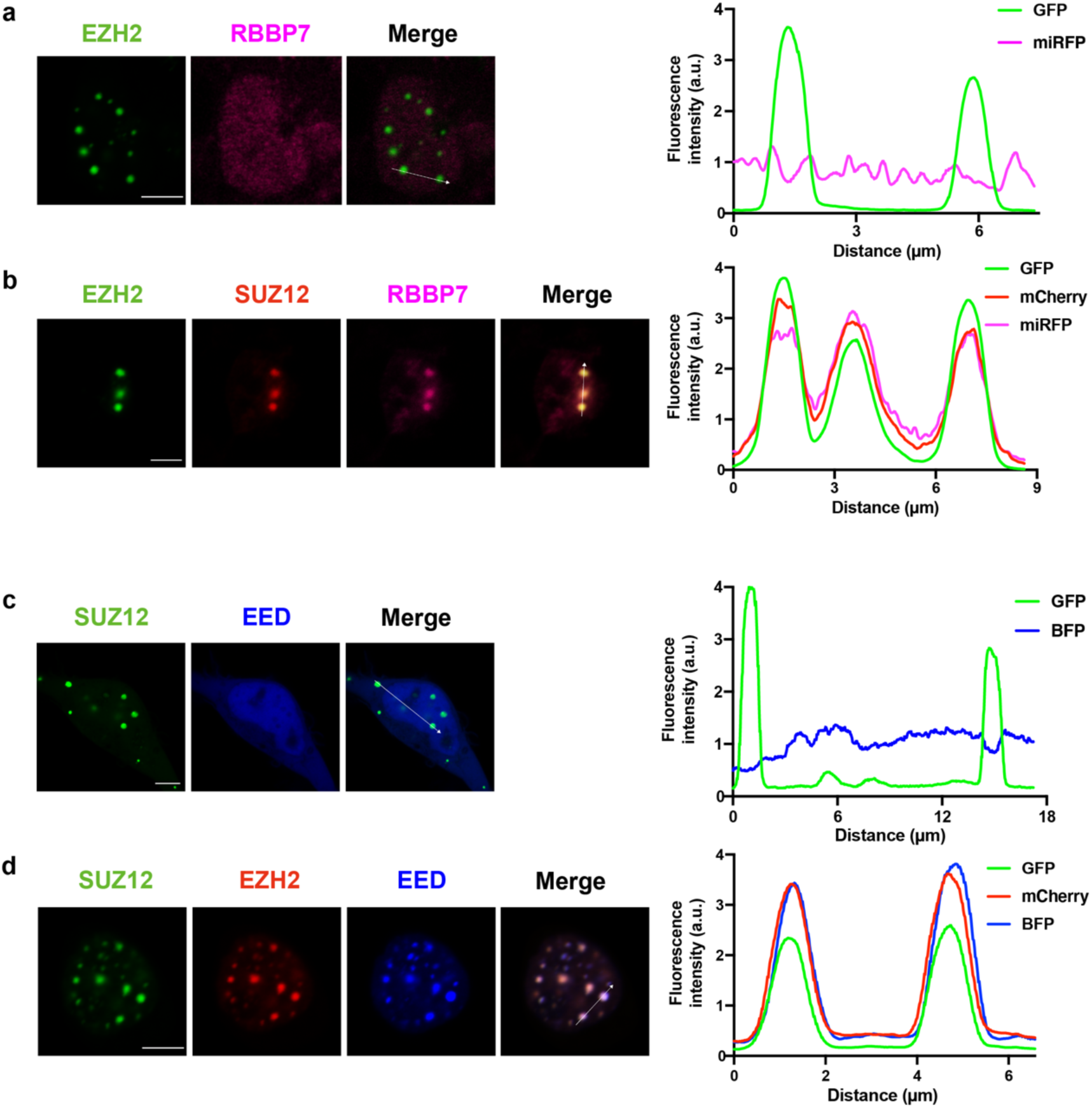
Compartmentalization of indirect protein-protein interactions in cells. **a**, Confocal images of HEK 293 cells overexpressing GFP-Nup98N-EZH2 and miRFP-RBBP7, with line analysis of fluorescence intensity on the right. The miRFP signal of the RBBP7 fusion protein remained evenly distributed in cells and was not enriched in the EZH2-containing compartments. The result was confirmed by plotting the fluorescence intensities along the line indicated by the white arrow in the merged confocal image. **b**, Confocal images of HEK 293 cells overexpressing GFP-Nup98N-EZH2, miRFP-RBBP7 and mCherry-SUZ12, with line analysis of fluorescence intensity on the right. The miRFP signal of RBBP7 and the mCherry signal of SUZ12 were enriched in the EZH2-containing compartments. The result was confirmed by plotting the fluorescence intensities along the line indicted by the white arrow in the merged confocal image. **c**, Confocal images of HEK 293 cells overexpressing GFP-Nup98N-SUZ12 and BFP-EED, with line analysis of fluorescence intensity on the right. The BFP signal of EED remained evenly distributed in cells and was not enriched in the SUZ12-containing compartments. The result was confirmed by plotting the fluorescence intensities along the line indicated by the white arrow in the merged confocal image. **d**, Confocal images of HEK 293 cells overexpressing GFP-Nup98N-SUZ12, BFP-EED and mCherry-EZH2, with line analysis of fluorescence intensity on the right. The BFP signal of EED and the mCherry signal of EZH2 were enriched in the SUZ12-containing compartments. The result was confirmed by plotting the fluorescence intensities along the line indicated by the white arrow in the merged confocal image. All scale bars, 5 μm.

## Discussion

Among numerous approaches to investigating PPIs in cells (Zhou et al., 2016), PACPIC stands out due to its simplicity and the balance between specificity and efficiency. Phase separation provides compartments for the enrichment of specific proteins, and hence greatly amplifies the signal-to-noise ratio. Recruitment into and enrichment within compartments require energy input to compensate the entropic penalty due to the concentration gradient. As such, only molecules with specific binding partners enriched within compartments can be effectively recruited, thus ensuring the specificity of PACPIC. The signals from protein interactions can be easily and faithfully detected within cells, without the requirement for time-consuming *in vitro* experiments such as protein purification, thus ensuring the efficiency of PACPIC.

Most importantly, in PACPIC, the recruitment, rather than the formation of phase separated compartments, serves as the readout of the interactions. This design protects the PPIs of interest from any artificial effects introduced by multivalency, as the client proteins are on separate vectors from that of the phase separation scaffold and thus do not undergo multimerization owing to phase separation.

In addition, the properties of liquid phases guarantee the relative freedom of movement of molecules and thus the reversibility of recruitment, making it possible to screen for inhibitors of specific PPIs and hence therapeutic drugs. Finally, as only a small number of cells is required for unambiguous readout of PPIs, PACPIC is also amenable to high-throughput screening (HTS) for modulators of certain PPIs that are already known for critical biological functions. It is worth noting that a similar method, PACPIT, which is suitable for HTS screening of PPIs *in vitro*, has been extensively evaluated (see the accompanying study for details).

This study is tailored for investigating direct and indirect interactions between proteins that are known and/or suspected to form a complex. By integrating proximity-labeling techniques into phase separation-derived compartments, our method can be extended to identify unknown binding partners of proteins.

## Acknowledgements

This work was supported by grants from the National Key R&D Program (2019YFA0508403 to P.L.), the National Natural Science Foundation of China (31871443 to P.L.; 31800637 to J.W.), and the Students Research Training Program of Tsinghua University (1911S0032 to H.L.).

## Author contributions

P.L. conceived and supervised the project; H.L. and J.W. carried out the experiments; H.L. and P.L wrote the manuscript.

## Materials and Methods

### Molecular Cloning

We cloned human the N-terminal half of Nup98 (1-515), full-length (FL) EZH1, FL EZH2, FL SUZ12 (1-739), truncated SUZ12 (561–739), FL EED, FL RBBP4 and FL RBBP7 into pCDNA3.1 expression vectors.

### Cell Culture

We maintained HEK 293 cells in DMEM medium (SH30022.01, Hyclone) with 10% fetal bovine serum (FBS), 100 μg/ml penicillin, 100 μg/ml streptomycin and 2 mM L-glutamine. For confocal microscope imaging, we seeded HEK 293 cells in a 4-chamber glass bottom dish (D35C4-20-1-N, Invitrogen). When the cells covered 70-80% of the chamber of a dish (almost 24 hours after seeding), we transfected them with plasmids (final concentration: 1 μg/ml) using Hifectin I (sj194, Applygen) as the transfection regent. After another 20-24 hours of culture, cells were imaged using a confocal microscope.

### Microscopy

We used a NIKON A1 microscope with 100 × oil immersion lens to image HEK 293 cells and NIS-Elements AR Analysis to analyze the images.

### Fluorescence recovery after photobleaching

We bleached cellular green fluorescence puncta with a 488 nm laser pulse (70% intensity; dwell time 0.5 s; n = 3), then we recorded the fluorescence intensity change with time. All the experiments were performed using a NIKON A1 microscope and data were analyzed by NIS-Elements AR Analysis.

### Data Analysis

We performed data analysis using Excel, NIS-Elements AR Analysis, and Prism8.

